# Artificial Intelligence Approaches to Assessing Primary Cilia

**DOI:** 10.1101/2021.02.03.429602

**Authors:** Ruchi Bansal, Staci E. Engle, Tisianna K. Kamba, Kathryn Brewer, Wesley R. Lewis, Nicolas F. Berbari

**Affiliations:** Department of Biology, Indiana University-Purdue University Indianapolis, Indianapolis IN, USA; Nikon Instruments Inc., New York, NY, USA; Stark Neurosciences Research Institute, Indiana University, Indianapolis IN, USA; Center for Diabetes and Metabolic Diseases, Indiana University School of Medicine, Indianapolis, IN, USA

## Abstract

Cilia are microtubule based cellular appendages that function as signaling centers for a diversity of signaling pathways in many mammalian cell types. Cilia length is highly conserved, tightly regulated, and varies between different cell types and tissues and has been implicated in directly impacting their signaling capacity. For example, cilia have been shown to alter their lengths in response to activation of ciliary G protein-coupled receptors. However, accurately and reproducibly measuring the lengths of numerous cilia is a time-consuming and labor-intensive procedure. Current approaches are also error and bias prone. Artificial intelligence (Ai) programs can be utilized to overcome many of these challenges due to capabilities that permit assimilation, manipulation, and optimization of extensive data sets. Here, we demonstrate that an Ai module can be trained to recognize cilia in images from both *in vivo* and *in vitro* samples. After using the trained Ai to identify cilia, we are able to design and rapidly utilize applications that analyze hundreds of cilia in a single sample for length, fluorescence intensity and colocalization. This unbiased approach increased our confidence and rigor when comparing samples from different primary neuronal preps *in vitro* as well as across different brain regions within an animal and between animals. Moreover, this technique can be used to reliably analyze cilia dynamics from any cell type and tissue in a high-throughput manner across multiple samples and treatment groups. Ultimately, Ai-based approaches will likely become standard as most fields move toward less biased and more reproducible approaches for image acquisition and analysis.

**SUMMARY:** The use of Artificial Intelligence (Ai) to analyze images is emerging as a powerful, less biased, and rapid approach compared with commonly used methods. Here we trained Ai to recognize a cellular organelle, primary cilia, and analyze properties such as length and staining intensity in a rigorous and reproducible manner.

## INTRODUCTION

Primary cilia are sensory organelles protruding from most mammalian cell types^1–4^. They are generally solitary appendages critical for coordinating diverse cell signaling pathways by integrating extracellular signals^5–7^. Primary cilia play important roles during embryonic development and adult tissue homeostasis, and disruption of their function or morphology is associated with several genetic disorders, which are collectively called ciliopathies. Due to the near ubiquitous nature of cilia, ciliopathies are associated with a wide range of clinical features that can impact all organ systems^8–12^. In animal models of ciliopathies, loss of ciliary-structure or signaling capacity manifests in several clinically relevant phenotypes including hyperphagia-associated obesity^3, 13–15^. In many model systems, cilia length changes have been shown to impact their signaling capacity and functions^16–19^. However, there are several time consuming and technical challenges associated with accurately and reproducibly assessing cilia length and composition.

The adult mammalian central nervous system (CNS) is one biological context that has posed a challenge for understanding cilia morphology and function. While it appears that neurons and cells throughout the CNS possess cilia, due to the limited tools and abilities to observe and analyze these cilia an understanding of their functions remains elusive^20^. For example, the prototypical cilia marker, acetylated α-tubulin, does not label neuronal cilia^20^. The difficulty of studying these cilia was partly resolved with the discovery of several G protein-coupled receptors (GPCR), signaling machinery and membrane associated proteins that are enriched on the membrane of neuronal cilia^21, 22^. All of these straightforward basic observations hint at the importance and diversity of CNS cilia, which thus far appears unparalleled by other tissues. For example, variation in cilia length and GPCR localization can be observed throughout the brain, with lengths in certain neuronal nuclei being different when compared with other nuclei^19, 23^. Similarly, their GPCR content and signaling machinery compliment show diversity based on neuroanatomical location and neuronal type^2, 24–29^. These simple observations demonstrate that mammalian CNS cilia length and composition are tightly regulated, just as in model organisms, like *Chlamydomonas reinharditii,* but the impact of these length differences on cilia function, signaling and ultimately behavior remains unclear^16, 30–32^.

Accurately measuring cilia length and composition proves to be a technical challenge prone to user error and irreproducibility. Currently cilia *in vivo* and *in vitro* are most often identified using immunofluorescent approaches that label ciliary proteins or cilia-enriched fluorescent reporter alleles^33–35^. The lengths of these fluorescently tagged cilia are then measured from a 2-dimensional (2D) image using line measurement tools in image analysis programs such as ImageJ^36^. This process is not only tedious and labor intensive but also prone to bias and error. These same obstacles arise when measuring cilia intensities, which help indicate changes in cilia structure^37^. To minimize the inconsistencies in these types of image analyses, artificial intelligence (Ai) programs are becoming more prevalent and affordable options^38^.

Ai is the advancement of computer systems that use the advantage of computer algorithms and programming to execute tasks that would usually require human intelligence^39^. Ai devices are taught to perceive recurring patterns, parameters, and characteristics and take actions to maximize the odds of creating successful outcomes. Ai is versatile and can be trained to recognize specific objects or structures of interest, such as cilia, and then be programmed to run a variety of analyses on the identified objects. Therefore, complex image data can be rapidly and reproducibly generated by Ai^38^. Automation and Ai analysis of captured images will increase efficacy and efficiency while limiting any potential human error and bias^39^. Establishing an Ai based methodology for cilia identification creates a consistent way for all research groups to analyze and interpret cilia data.

Here we utilize an Ai module to identify cilia both *in vivo* and *in vitro* on 2D images. Using a set of sample images the Ai is trained to identify cilia. Once training is complete, the designated Ai is used to apply a binary mask over Ai identified cilia in an image. The binaries applied by the Ai are modifiable, if necessary, to ensure all cilia in the images are properly identified and non-specific identification is eliminated. After utilizing the Ai to identify cilia, custom-built general analysis (GA) programs are used to perform different analyses such as measuring cilia length and fluorescence intensity. The data collected is exported into a table that can be easily read, interpreted, and used for statistical analyses (**Fig 1**). The use of automated technology and Ai to identify cilia and obtain specific measurements between experimental groups will aid in future studies aimed to understand the impact of CNS cilia function and morphology on cell-cell communication and behavior.

**Figure 1.**
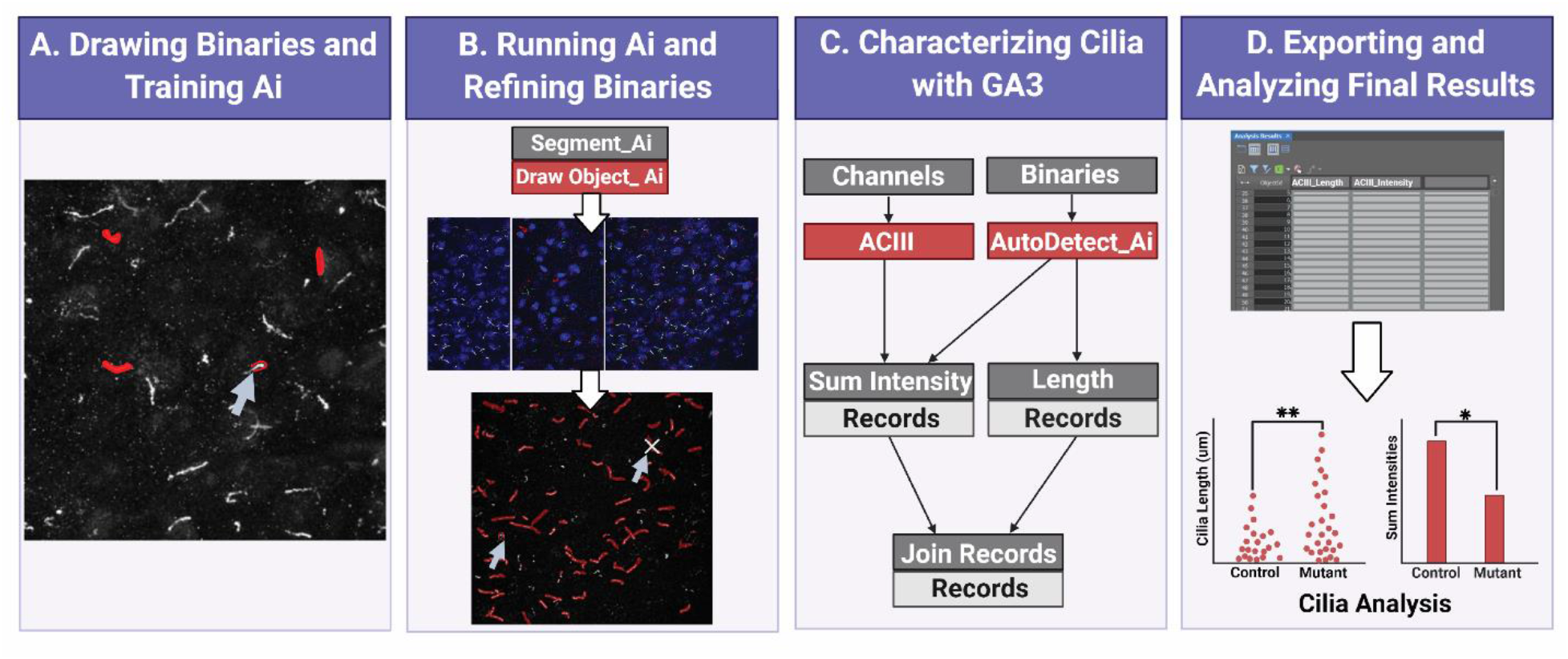
Workflow for measuring cilia length and intensity using Ai. **A.** To train the Ai, binaries are drawn around the object of interest (cilia) on the raw training images. Using the drawn binaries, Segment Ai is trained to recognize the shape and pixel intensities of the cilia. **B.** Next, the trained Segment Ai is applied to raw experimental images. It draws binaries on objects it recognizes as cilia. These binaries can be refined to makes sure all and only cilia are being analyzed. **C.** A GA3 program is constructed to analyze the intensity and length of objects recognized by the Ai. **D.** The records are imported into a table in the software. This table can then be exported for further analysis.

## PROTOCOL

1. <u>Acquire raw images</u>
  1.1. Fix and immunolabel samples as needed^20^.

1.1.1 Image cilia using a confocal microscope at maximum bit depth using the same pixel size with Nyquist resolution.
1.1.2 Export images as monochrome Tagged Image Format (.tif) files.
  NOTE: This protocol outlines how to use the Ai module specifically within NIS Elements software. If images were acquired as .nd2 files, exporting images as .tif files is not necessary and the user can proceed directly to step 2.3. If images were acquired on a different system, a NIS Elements license can be purchased separately and .tif files can be converted as outlined in the next steps.
2. <u>Train Ai to identify cilia</u>
  2.1. Open training dataset

2.1.1. Select approximately 50 sample images with at least one cilia per frame to train the software and copy them in a single folder. This folder is used to direct the software when opening the images. Open these 50 frames within a single ND2 document with at least one cilia per frame. Pathway in video: (*File > Import/Export > Create ND File from File Sequence*).
2.1.2. Select the folder containing the training dataset. This will open the list of files in the center of the dialogue window. Define manually the organization of the files using at least one option in the drop-down menu above. The options are multipoint (for multiple maximum projection files), Z series (for a z stack image), time (for a time-lapse image), and wavelength (for files from multiple channels).
2.1.3. Enter the corresponding numerical values under each selected option. Select *‘None’* wherever options are not selected. Click *‘Convert’* to open the ND document.
  2.2. Calibrate images

2.2.1. Enter the pixel size in the lower left-hand corner of the image. Pathway in video: (*Right click uncalibrated > calibrate document > pixel size*)
  2.3. Identify cilia

2.3.1. Hand-identify the cilia by precisely tracing individual ciliary structures on all the opened frames using either *AutoDetect* or *Draw Object* in the *Binary Toolbar*. This will draw binary masks on the objects of interest. These binaries will serve as sample objects for training the software to identify cilia on pixel-based characteristics in future experimental image analysis. Pathway in video: (*View > Analysis Controls > Binary Toolbar > Draw Object*). NOTE: Remove any frame that does not have any binaries as the software will not begin training unless it is able to detect binaries on all of the opened frames.
  2.4. Train Ai

2.4.1. Begin training the software. This will open the train Segment.ai box. Pathway in video: (*NIS.ai > Train Segment.ai*).
2.4.2. In the ‘Train Segment.ai’ box, select the source channel to be used for training. If files from multiple channels are open, select only one channel as the source channel. Then select the appropriate ground truth binaries on which to train the Ai. Finally select the number of iterations needed to train the Ai depending on the size and distribution of the binaries. NOTE: If the binaries are easily detectable from the surroundings and well distributed throughout the image, the software may need less than 1000 iterations to get trained to identify the images. If the images have a low signal-to-noise ratio, it is ideal to run at least 1000 iterations while training to enable the Ai to identify cilia in test samples with high confidence.
2.4.3. Select the destination folder to save the trained Ai file (.sai) and *click ‘Trair’* to train the software. The software will now proceed to train itself to identify cilia based on the traced binaries. This process takes several hours. NOTE: When training, the software will display a graph showing training loss. The graph will initially show a spike before tapering to ideally around 1% loss where it plateaus for the rest of the training. Save the graph for future reference by checking the box for *‘Save graph screenshot’* on the Train Segment.ai box (**Supplemental Fig 1**).
2.4.4. If further refinement to the training is required, then continue training on the same dataset. Alternatively, train on a new dataset with the exact same parameters. It is not advisable to train an already trained Ai on a new dataset with different parameters or different objects of interest. Pathway in video: (*TrainSegment.ai > Continue training on* > select trained Ai file).
3. <u>Identify cilia using trained Ai</u>
  3.1. Open experimental dataset

3.1.1. Open the experimental confocal images of cilia in the software by converting the sample .tif files to .nd2 files, similar to step 2.1. Pathway in video: (*File > Import/Export > Create ND File from File Sequence*) NOTE: The images should be of the same pixel size as those used for training the Ai. If images are already in ND2 format, skip to step 3.3.
  3.2. Calibrate images

3.2.1. Enter the pixel size in the lower left-hand corner of the image. Pathway in video: (*Right click uncalibrated > calibrate document > pixel size*)
  3.3. Run the trained Ai on the opened files

3.3.1. Begin identifying the cilia using Ai. The software will now draw binaries on the cilia based on the training it received in the prior step. This process will take a few seconds. Pathway in video: (*NIS.ai > Segment.ai*). NOTE: The software will prompt to select the channel if multiple channels are open. Channels are listed by their respective names here. If not, the box marked ‘Mono’ will be automatically selected.
  3.4. Check images for misidentified binaries

3.4.1. Once the Ai has identified cilia and drawn binaries, check the images for any mistakenly identified object. If desired, manually delete any misidentified binaries. Pathway in video: (*View > Analysis Controls > Binary Toolbar > Delete Object*)
4. <u>Measuring cilia length and intensity</u>
  4.1. Create new General Analysis 3 (GA3) recipe

4.1.1. Now that the cilia have been identified and segmented, proceed to analyze different parameters of cilia such as lengths and intensities using GA3 tool. This will open a new window with a blank space in the center where the analysis will be defined. Pathway in video: (*Image > New GA3 Recipe*).
  4.2. Select the binaries to analyze

4.2.1. Since the cilia are already segmented using Segment.ai, GA3 will automatically detect the binaries appropriately labelled according to the Ai and include the node *‘Binaries* > *Auto Detect_AI’* or *‘Binaries* > *Draw Object_AI’.*
  4.3. Select the channels required for analysis

4.3.1. GA3 will also automatically detect the channels in the images and display their tabs under *‘Channels’.*
  4.4. Remove objects touching the border of the frame

4.4.1. Since the Ai will segment all cilia like objects in the frame, it will also detect incomplete cilia along the edges of the frame. These objects can either be removed manually in step 3.4 or they can be removed automatically in GA3. Pathway in video: (*Binary processing > Remove objects > Touching Borders)*
  4.5. Select parameters to measure cilia

4.5.1. Drag and drop the parameters to measure such as cilia length (*Length*) and intensities (*Sum Object Intensity*). Connect the nodes to the appropriate binary node (connection A) and channel nodes (connection B). Hover over the node connection for a tooltip to show which connection the node belongs to. Pathways in video: (*Measurement > Object Size > Length)* and (*Measurement > Object Intensity > Sum Obj Intensity)* NOTE: The node *‘Length’* connects only to the binary node, whereas *‘Sum Obj Intensity’* connects to both the binary and the channel nodes.
  4.6. Append the measurements in a single table

4.6.1. Combine all the measurements in a single output table by dragging and dropping the node *‘Append Column’* the analysis flow chart and connect it to the measurement nodes, *‘Length’* and *‘Sum Obj Intensity’*. Pathway in video: (*Data Management > Basic > Append Column)*
  4.7. Measure cilia

4.7.1. Measure cilia by clicking *‘Run’.* This process takes a few moments to measure all the cilia in the experimental images. The lengths and intensities will appear in a new *Analysis Results* window. NOTE: The table can sometimes include data from masks that Ai recognized as cilia but were too small to be detected by the human eye and eliminated in step 3.4. These objects can be removed from the data set by using a filter before statistical analysis. Here, a filter of 1 μm was used for *in vitro* cilia length measurements in **Figure 2** and 2 μm for *in vivo* cilia. This can be done prior to exporting data using the pathway below. Pathway in video: (*Analysis Results window > Define filter > Enter Value > Use Filter)*
  4.8. Export data

4.8.1. Export data for statistical analysis.
5. <u>Colocalization studies</u> NOTE: Colocalization analysis can be included in the same GA3 recipe used for measurements of cilia length and intensity analysis. If using the same recipe, open files as described below and measure the lengths and intensities of both channels along with the colocalization coefficients in the same analysis pipeline.
  5.1. Open experimental dataset

5.1.1. Open the experimental confocal images of cilia in the software by converting the sample .tif files to .nd2 files Pathway in video: (*File > Import/Export > Create ND File from File Sequence*).
5.1.2. In the pop-up window, select your 16-bit depth monochrome files from all channels of interest from the window explorer located in the first column of the pop-up window. Select *“Multipoint”* or “*Z Series”* from the first drop down menu and enter a value corresponding to the total number of images or stacks, respectively.
5.1.3. In the second drop down box, select “Wavelength” and change the value to the total number of channels in the folder. The software will automatically unlock a *“Wavelength”* selection window located at the bottom right end of the pop-up window. Use the *“Color”* drop down menu to select the color of each channel. Provide each channel with a different name under the *Name* column. Once all information is updated, click *‘Convert’.* The software will automatically generate an *“All”* image file overlayed with all the individual images from all the selected channels.
  5.2. Calibrate images

5.2.1. Enter the pixel size in the lower left-hand corner of the image. Pathway in video: (*Right click uncalibrated > calibrate document > pixel size*)
  5.3. Run the trained Ai on the first channel

5.3.1. Begin identifying the cilia on one of the opened channels (eg. ACIII; **Fig 5A**) using Ai. The software will now draw binaries on ACIII labelled cilia based on the training it received for this channel. This process will take a few seconds. Pathway in video: (*NIS.ai > Segment.ai > Source channels > ACIII*).
  5.4. Run the trained Ai on the second channel

5.4.1. Begin identifying the cilia on the other opened channel (eg. MCHR1; **Fig 5B**) using Ai. The software will now draw binaries on MCHR1 labelled cilia based on the training it received for this channel. This process will take a few moments. Pathway in video: (*NIS.ai > Segment.ai > Source channels > MCHR1*).
  5.5. Check images for misidentified binaries

5.5.1. Once the Ai has identified cilia and drawn binaries, check the images for any misidentified objects. Manually delete any misidentified binaries if necessary. Pathway in video: (*View > Analysis Controls > Binary Toolbar > Delete Object)*
  5.6. Create new GA3 recipe

5.6.1. Now that the cilia have been identified and segmented, proceed to the colocalization analysis using the GA3 tool. This will open a new window with a blank space in the center where the analysis will be defined. A window with all identified binaries and channels will be generated. Verify that all desired channels and binaries required for analysis are present and selected. Pathway in video: (*Image > New GA3 Recipe*).
  5.7. Remove objects touching the border of the frame.

5.7.1. Since the Ai will segment all cilia-like objects in the frame, it will also detect incomplete cilia along the edges of the frame. These objects can either be removed manually in step 5.5 or they can be removed automatically in GA3. Pathway in video: (*Binary processing > Remove objects > Touching Borders)*
  5.8. Setup the colocalization pathway in GA3

5.8.1. To measure overlap of the two channels within cilia, use Mander’s Coefficient Correlation. Drag and drop the *Manders Coefficient* node into the blank space of the GA3 recipe and connect it to the appropriate binary and channels. Here, ‘connection A’ connects with the ACIII binary, ‘connection B’ with the MCHR1 channel and ‘connection C’ with the ACIII channel to determine the overlap of MCHR1 within ACIII binary. Pathway in the video: (*Measurement > Object Ratiometry > Manders Coefficient)* NOTE: The software allows for measuring colocalization using Pearson Coefficient Correlation using the same steps as described in this protocol^40^.
  5.9. Append the measurements in a single table

5.9.1. Combine all the measurements in a single output table. Pathway in video: (*Data Management > Basic > Append Column)*
  5.10. Measure colocalization

5.10.1. Measure cilia by clicking *‘Run’.* This process takes a few moments to measure all the cilia in the experimental images. The data will appear in a new *Analysis Results* window.
  5.11. Export data

5.11.1. Export data for statistical analysis.

**Figure 2.**
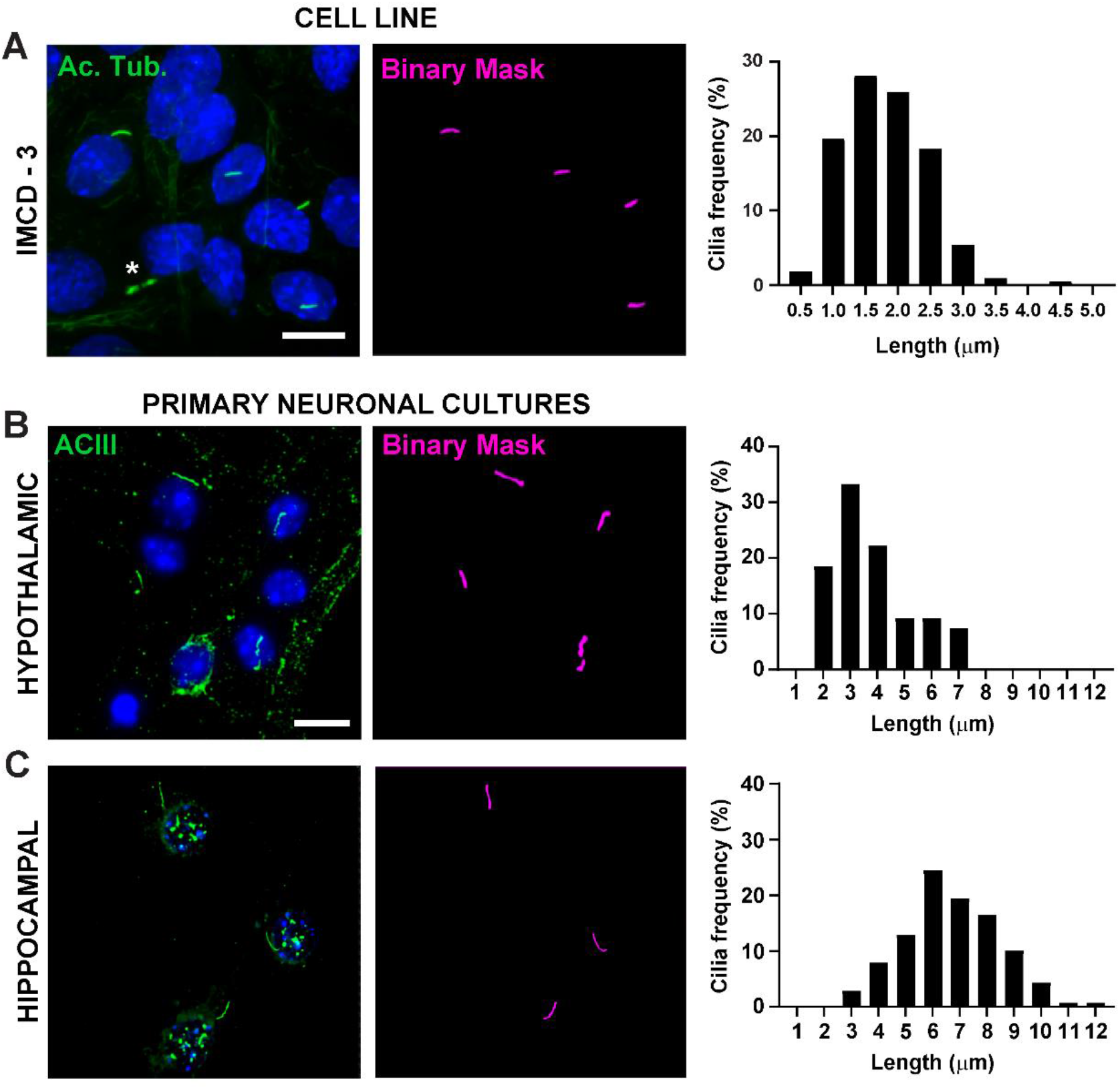
*In vitro* cilia length measurements. Representative images of cilia in **A.** IMCD cells (green, acetylated tubulin), **B.** primary hypothalamic cultures (green, ACIII) and **C.** hippocampal cultures (green, ACIII). A trained Ai was used to recognize cilia as shown in the binary mask (magenta) and then GA3 was used to measure cilia length. Distribution of cilia length is graphed as percentage of cilia in 0.5 or 1.0-micron bins. * indicates cytokinetic bridge properly not recognized by Ai. n=225 cilia in IMCD cells from 3 replicates, 54 cilia in hypothalamic and 139 cilia in hippocampal cultures from 3 animals. Scale bars 10 μm.

## RESULTS

### Training Ai to identify cilia

Measuring and assessing cilia structural length and composition can be a tedious, time consuming, and error prone process. Here, we use Ai for accurately segmenting cilia from a large pool of images and analyze their lengths and intensities with an analysis tool **(Fig 1)**. All Ai approaches require training steps for their implementation. We established a training pipeline to recognize cilia, which was performed by manually applying binary masks on ciliary structures. This information is then used to train the Ai based on pixel characteristics under the applied binaries. As a general guideline, the training involves the software going through several iterations, approximately 1000, and is considered optimal if the training loss or error rate is less than 1%. However, the number of iterations and errors in the training process can vary depending on the sample images used for training. For example, after our training sessions using *in vitro* neuronal cilia images, the error rate was 1.378% compared with 3.36% for *in vivo* brain section images (**Supplemental Fig 1**). Once training is completed, Ai can then be used to segment cilia from experimental images within seconds and the resultant binary masks are used for measuring structural parameters. This eliminates the need to segment objects using the traditional method of intensity thresholding which may be difficult in images with high background noise or when objects are in close proximity to each other. Ai also reduces the potential for error and bias by applying the same algorithm across all images, regardless of user.

### Measuring cilia length using GA3

Cilia length is tightly regulated and is associated with functional impacts on ciliary signaling^16, 19^. Here, we measured cilia lengths using an analysis pipeline within NIS Elements software called General Analysis 3 or GA3. GA3 aids in combining multiple tools in a single workflow to build customized routines for each experiment. We began by measuring cilia lengths in a cell line. Cilia on mouse inner medullary collecting duct (IMCD-3) cells were immunolabelled with acetylated tubulin and imaged using a confocal microscope. We measured cilia lengths using GA3 after segmenting with segment.ai **(Supplementary Fig 3A)**. While acetylated α-tubulin is preferentially found in the primary cilium, it is also found in other microtubule rich regions such as the cytoskeleton as well as the cytokinetic bridge. The trained Ai properly identified cilia in the image but not other non-ciliary, acetylated tubulin positive structures. Cilia on IMCD cells ranged from 0.5 μm to 4.5 μm with an average length of 1.8 ± 0.04 μm **(Fig 2A)**. We next tested the ability of Ai to measure cilia lengths in primary neuronal cultures. We cultured neurons from the hypothalamus and hippocampus of neonatal mice for 10 days and immunolabelled them with the cilia marker adenylate cyclase III (ACIII)^41, 21^. When analyzing neuronal cultures, we found it useful to apply a filter prior to statistically analyzing the lengths. Because of a higher signal to noise ratio, several objects less than 1 μm were identified that were not cilia. Therefore, we filtered the data to eliminate any objects that were less than 1 μm in length to ensure that only cilia were analyzed. In cultured hypothalamic neurons, cilia lengths ranged from 2 μm to 7 μm with an average length of 3.8 ± 0.19 μm **(Fig 2B)**. Interestingly, cultured hippocampal neuronal cilia were longer with a mean length of 6.73 ± 0.15 μm **(Fig 2C)**. It has been reported that different neuronal nuclei within the hypothalamus show distinct cilia lengths and that these cilia alter their lengths in response to physiological changes in a nucleus specific manner^19, 23^. Therefore, we also labelled hypothalamic brain sections from adult male C57BL/6J mice with ACIII and imaged the arcuate nucleus (ARC) and paraventricular nucleus (PVN). Using GA3 to measure cilia lengths, we observed that the *in vivo* hypothalamic cilia appeared longer than *in vitro* cilia. Specifically, hypothalamic cilia *in vivo* range from 1 μm to about 15 μm **(Fig 3)**. There were no significant differences between the lengths of cilia in the PVN (5.54 ± 0.0.42 μm) and those in the ARC (6.16 ± 0.27 μm) **(Fig 3C)**^23^. Similarly, cilia in the cornu ammonis (CA1) region of the hippocampus display a narrower length range from 1 μm to 1 0 μm with an average length of 5.28 ± 0.33 μm **(Fig 3)**. In accordance with previously published studies, our analysis using Ai and GA3 tools showed that cilia from different brain regions show diversity in length^19, 23^. Moreover, using this Ai approach we are able to rapidly assess a large number of cilia.

**Figure 3.**
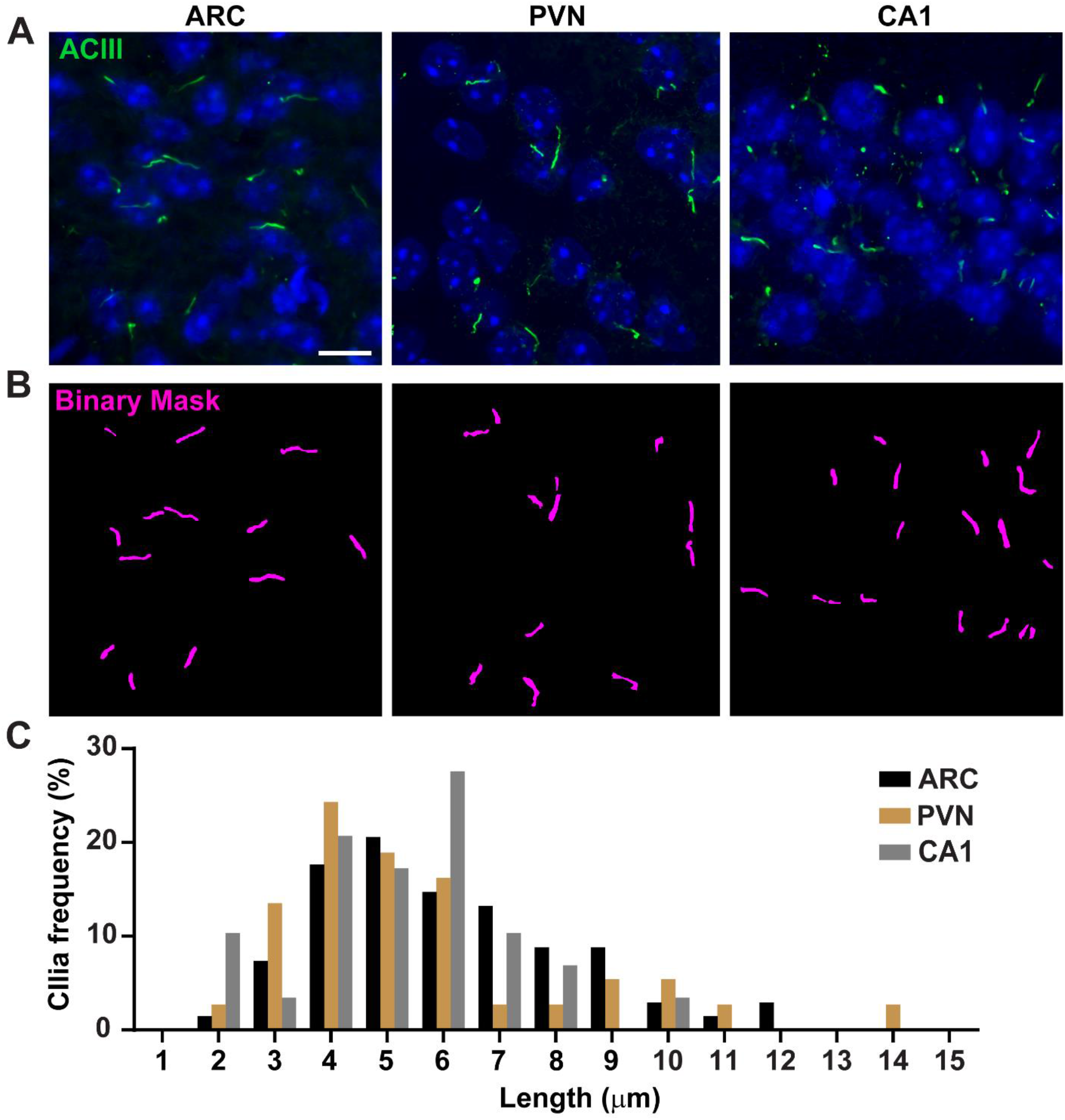
*In vivo* cilia length measurements. **A.** Representative images of cilia (green, ACIII) in the ARC, PVN, and CA1 of adult mouse brain sections. **B.** A trained Ai in NIS Elements was used to recognize cilia as shown in the binary mask (magenta) and then GA3 was used to measure cilia length. **C.** Distribution of cilia length is graphed as percentage of cilia in one-micron bins. n= 68 cilia in ARC, 36 in PVN and 29 in CA1 from 3 animals. Scale bars 10 μm.

### Measuring cilia composition using GA3

The primary cilium is a signaling hub for many pathways which utilize diverse types of proteins to carry out unique functions such as motor proteins, intraflagellar transport proteins and GPCRs to name a few^3, 24, 42, 43^. Maintaining the appropriate levels of these proteins within the cilium is important for proper functioning and often appears cell context dependent. Fluorescent labelling of these proteins has not only enabled us to visualize them but also quantify their intensities as a measure of the amount of the labelled protein within the relatively small compartment^20^. Therefore, we sought to determine the intensities of a ciliary GPCR, Melanin Concentrating Hormone Receptor-1 (MCHR1), *in vivo* in both the ARC and PVN of the hypothalamus of adult male mice^24, 44^. Using Ai and GA3, we measured the lengths of MCHR1 positive cilia along with intensities to ensure the objects being counted were cilia **(Supplementary Fig 3A)**. We eliminated objects post-analysis that were less than 2 μm in length and analyzed intensities of remaining binary masks. Interestingly, we found that the intensity of ciliary MCHR1 in PVN is significantly higher than that in ARC indicating a stronger presence of ciliary MCHR1 in PVN **(Fig 4)**. Further studies are required to determine the significance of ciliary MCHR1 in these neuronal circuits. We also measured the intensities of ciliary MCHR1 in primary cultured neurons of hypothalamus and hippocampus. Cilia from both cultures display a wide distribution of MCHR1 intensities suggesting the presence of heterogenous neuronal populations **(Supplementary Fig 2)**. Thus, using sophisticated analytical tools like Ai and GA3 allows for the evaluation of cilia heterogeneity within the same tissue or between multiple tissues. It will be interesting to see if other neuronal GPCRs show similar differences in their localization within neurons of the same tissue and if this alters in response to physiological changes.

**Figure 4.**
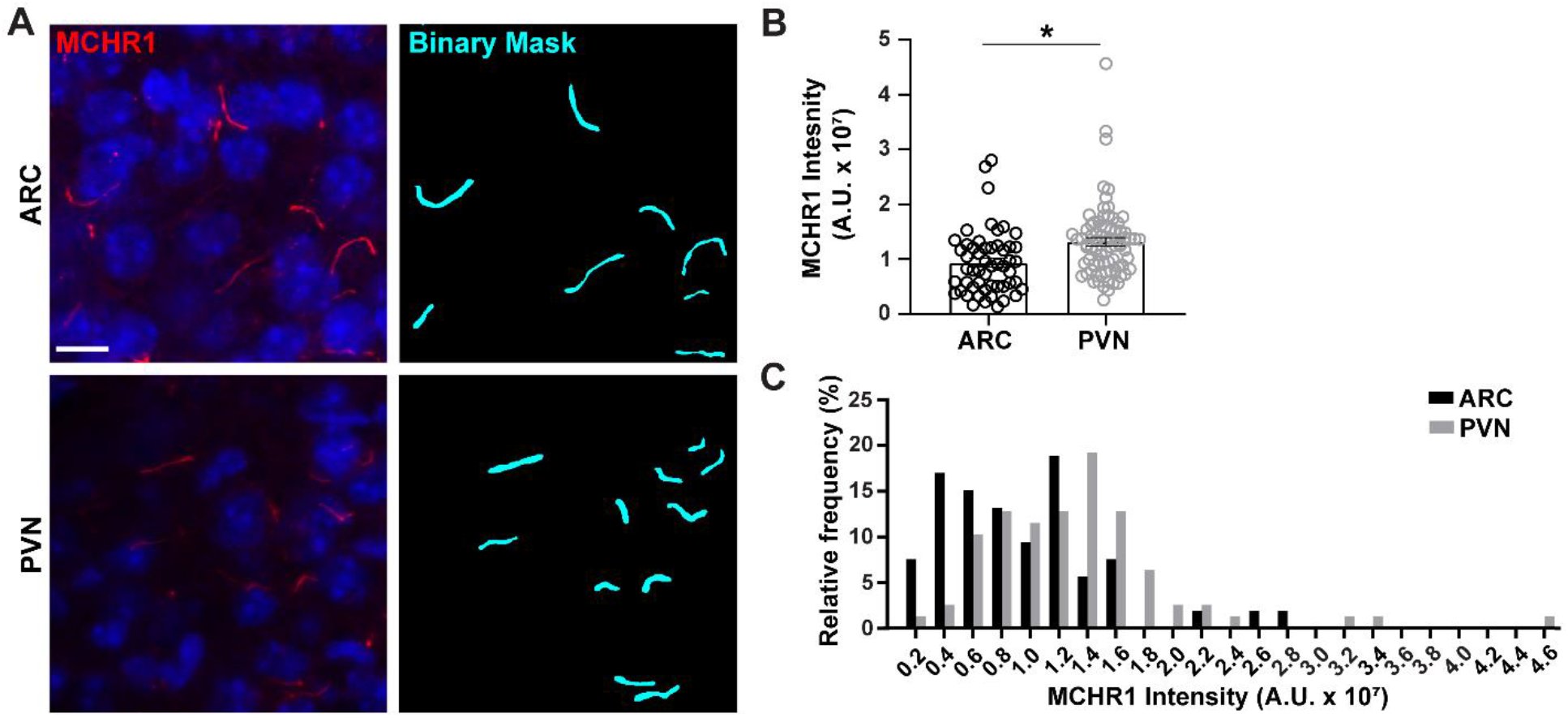
Ai assisted cilia staining intensity measurements of hypothalamic neuronal cilia. **A.** Representative images of cilia (MCHR1, red) in the ARC and PVN, of adult mouse brain sections. A trained Ai in NIS Elements was used to recognize cilia as shown in the binary mask (cyan) and then GA3 was used to measure the intensity of MCHR1 staining in cilia. **B.** MCHR1 intensities are graphed as average ± S.E.M. Each dot represents a cilium. * p < 0.05, Student’s t-test. **C.** Distribution of MCHR1 intensity is graphed as percentage of cilia in bins of 0.2 x 10^7^ Arbitrary Units (A. U.). n= 53 cilia in ARC, 78 in PVN from 3 animals. Scale bars 10 μm.

**Figure 5.**
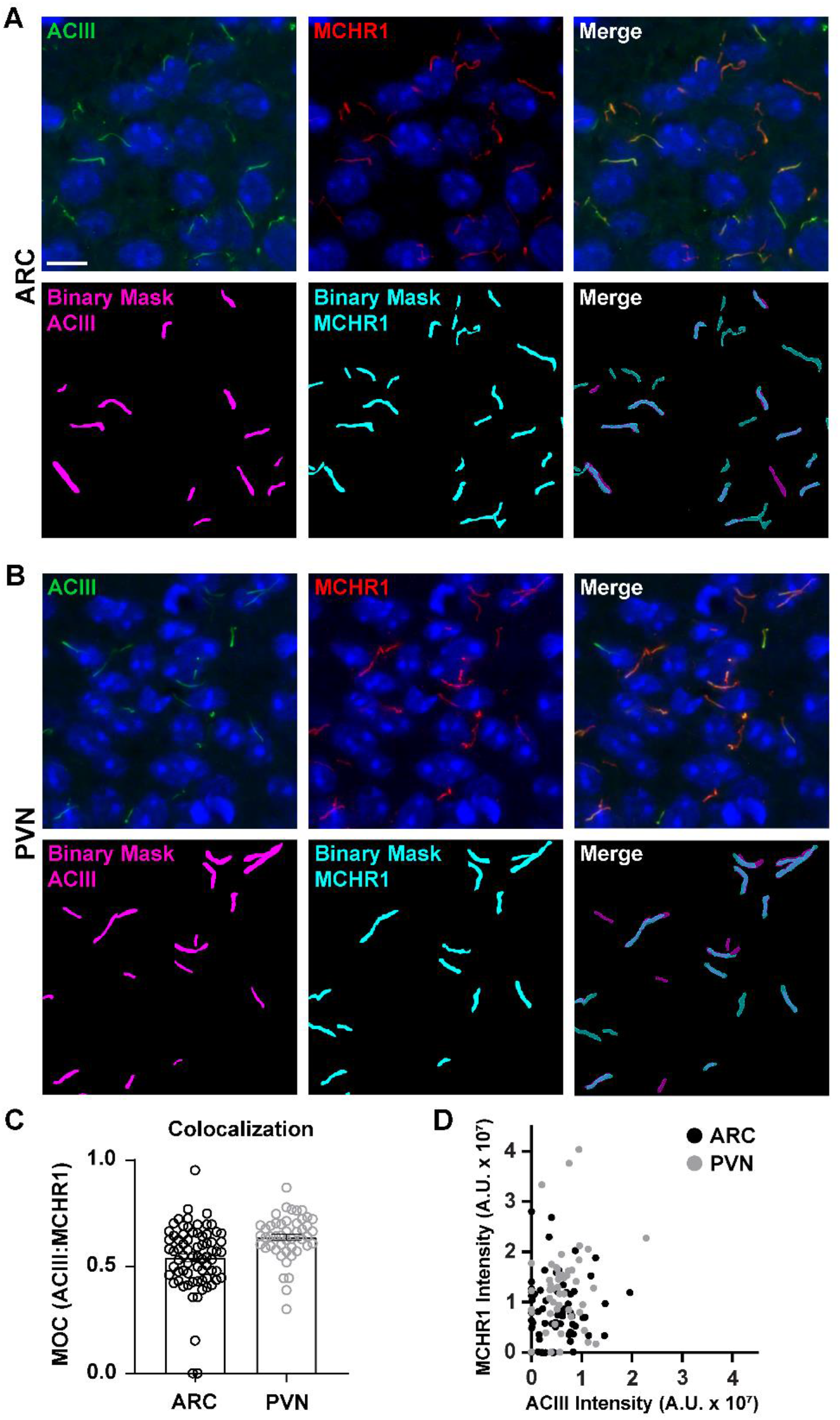
Ai assisted cilia colocalization analysis. **A.** and **B.** Representative images of cilia in the ARC and PVN respectively. Cilia are labelled with ACIII (green) and MCHR1 (red). **B.** A trained Ai in NIS Elements was used to recognize cilia as shown in the binary mask (magenta for ACIII labelled cilia, cyan for MCHR1 labelled cilia). GA3 was used to recognize cilia that contained both ACIII and MCHR1. **C.** Manders overlap coefficient (MOC) values are graphed as average ± S.E.M. Each dot represents a cilium. * p < 0.05, Student’s t-test. **D.** Scatter plot of MCHR1 intensity vs. ACIII intensity in ARC and PVN. Each dot represents a cilium. n= 72 cilia in ARC, 47 in PVN from 3 animals. Scale bars 10 μm.

### Colocalization

While measuring fluorescence intensities within a complete image field can give an impression of protein, it fails to provide information such as spatial distribution or proximity to other nearby proteins and cellular structures. Here, we measured the overlap of MCHR1 with ACIII as a cilia marker by plotting the intensities of MCHR1 against that of ACIII for each binary mask **(Fig 5)**. The graph shows that majority of the cilia are positive for both, ACIII and MCHR1, though some cilia show stronger expression of one channel over the other. Furthermore, there are some cilia that show the presence of either ACIII or MCHR1 as is evident from the points that lie directly on the x-axis and y-axis respectively. To quantify this overlap, we measured Mander’s overlap coefficient and compared the extent of MCHR1 expression in neuronal cilia of the ARC and PVN^40^. Interestingly, our analysis revealed that there was a significant increase in the coefficients of the PVN (0.6382 ± 0.0151) than those in the ARC (0.5430 ± 0.0181) **(Fig 5C)**. This is consistent with our previous data where we observed higher MCHR1 intensities in PVN compared to ARC **(Fig 4)**. This data suggests that like cilia length, the expression pattern of MCHR1 in the ciliary compartment varies in different regions of the brain. Using the same analysis pipeline, it will be possible to determine whether other ciliary GPCRs such as Neuropeptide Y Receptor Type 2 (NPY2R) and Somatostatin Receptor Type 3 (SSTR3) show similar amounts of diversity.

### Measuring intensity profile along the cilium

Once cilia have been identified using segment.ai, the GA3 recipe can be modified to combine cilia analysis with identification of other structures of interest in the image. For example, labelling with basal body markers is useful for identification of cilia polarity. To do this analysis, we imaged hypothalamic brain sections from P0 mice that express ARL13B-mCherry and Centrin2-GFP and imaged the arcuate nucleus (ARC) and paraventricular nucleus (PVN)^34^. Here, cilia were identified using Ai as before but now the modified GA3 recipe includes identification of Centrin2-GFP, a centriolar protein found at the base of cilia (**Supplementary Fig 3B**). By labelling Centrin2-GFP, the base of cilia can be distinguished from the tips of ARL13B-mCherry positive cilia (**Fig 6A**). Then, instead of measuring intensity within the whole cilia, we are able to measure changes in ARL13B intensity along the length of cilia (**Fig 6B**). We can also compare differences in the intensity of ARL13B between the proximal ends and distal ends of the cilia. To do this, we divided the cilium length in 1-micron bins starting from the base and designated the first micron bin as the proximal end and the last micron bin as the distal end. Our analysis revealed that there is significantly more ARL13B present closer to the base than the tip of the cilium in both ARC and PVN, and this is consistent with previously published studies in human chondrocytes^45^ (**Fig 6C**). In this type of analysis, instead of applying a length filter to exclude small non-ciliary objects from analysis, only cilia associated with Centrin2-GFP labelling are analyzed. This can be advantageous in situations where genetic mutations render very short cilia, or if changes in cilia subdomains like the transition zone or tip have been implicated. Identification of cilia using Ai and GA3 analysis is highly adaptable and can be tailored to fit a variety of complex research questions.

**Figure 6.**
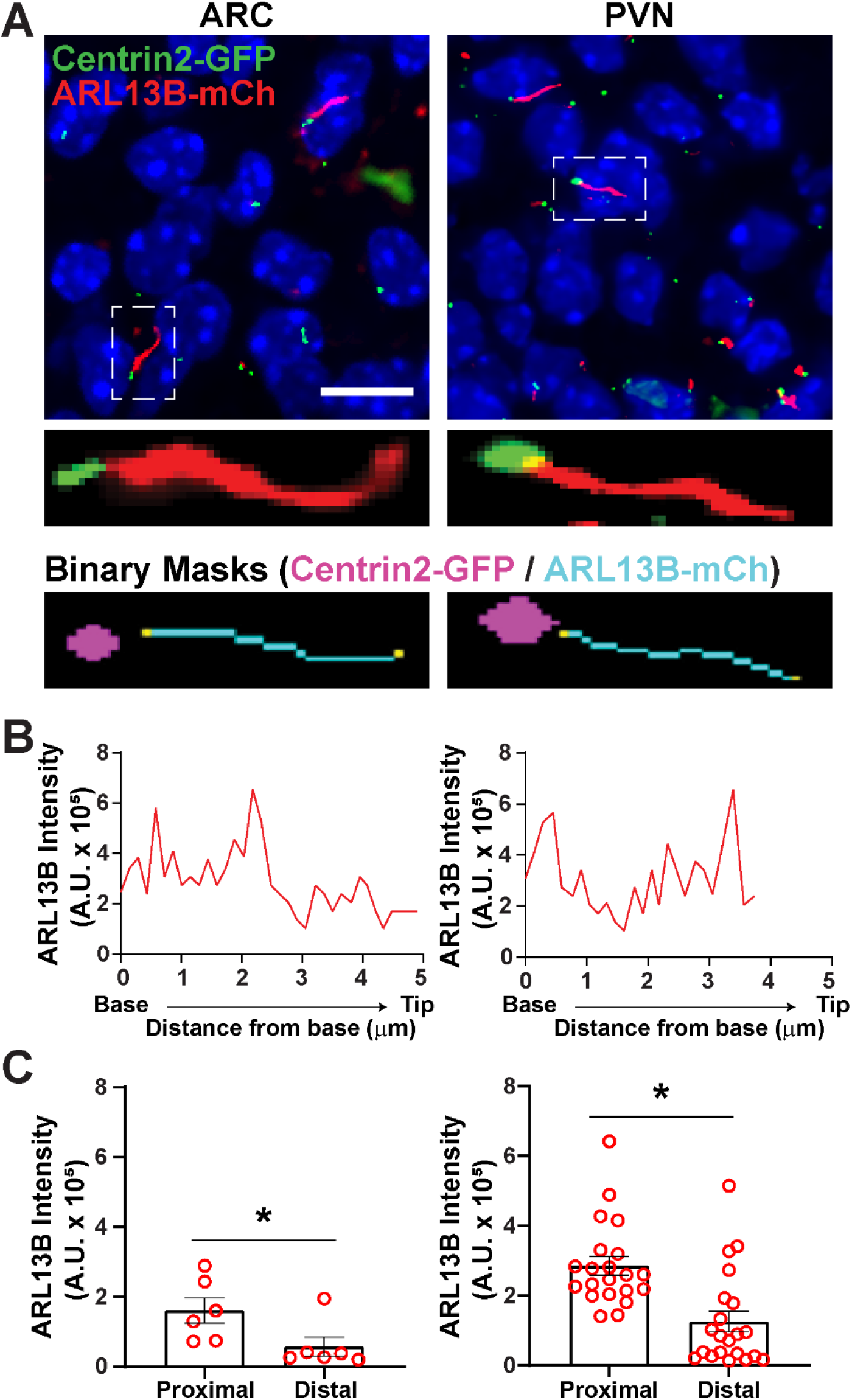
Cilia and basal body analysis. **A.** Representative images of cilia (red, ARL13B-mCherry) and basal body marker (green, Centrin2-GFP) in the ARC and PVN of P0 mice. A trained Ai was used to recognize cilia as shown in the binary mask (cyan). The binary mask for basal body (magenta) was drawn by thresholding in GA3 recipe. **B.** Representative line scan intensity of a cilium. **C.** ARL13B intensities at the proximal and distal ends of Ai identified cilia graphed as average ± S.E.M. The proximal and distal ends are defined as the region within the first 1 μm length and the last 1 μm length respectively from the base of the cilium. Each dot represents a cilium. * p < 0.05. n = 6 cilia in ARC from 2 animals and 21 cilia in PVN from 3 animals. Scale bars 10 μm.

## DISCUSSION

Length and intensity measurements are common ways primary cilia are analyzed, however, there is not a standardized conventional method used in the field. Identification and quantification of primary cilia using software such as ImageJ is time consuming and prone to user bias and error. This makes it difficult to accurately analyze large data sets. Here we show that using an Ai program can overcome many of these challenges making high throughput analysis of primary cilia achievable. Herein we describe the procedure for training an Ai-based application to recognize primary cilia and outline the steps required to analyze length and intensity.

While the initial training of the Ai to recognize cilia requires significant time from the user, once completed it can be used on any data set acquired with the same parameters. The binary mask generated by the Ai is modifiable such that any errors can be corrected. However, errors in cilia identification should signal the user that the Ai needs to be further trained with additional images. One major advantage of this method is that the Ai can be trained to recognize cilia in different sample types in both 2D and 3D. Previous analysis methods generated within labs have various limitations including requiring manual thresholding for identification and problems identifying cilia imaged from tissue sections where the cell density is high^36,46,47^. These methods are also specialized for cilia analysis whereas analysis using NIS Elements software can evaluate several aspects of the images simultaneously. Because the Ai described here is part of the NIS Elements software package, images acquired using a Nikon microscope can be easily continued through to analysis. However, imaging with Nikon is not required for use of this this method. Regardless of the captured raw data file format, “.tif” files can be opened by NIS Elements to use in the Ai.

This Ai application within NIS Elements is widely available and possibly already part of image analysis software in use by labs studying primary cilia. With the prevalence of Ai technology expanding, other imaging software may expand their analysis options to include a similar Ai module. Applying Ai analysis to cilia identification can be used for several different aspects of cilia analysis. While we outlined methods for a few simple analyses such as length **(Fig 2 and 3)**, intensity **(Fig 4)** and colocalization **(Fig 5)** more sophisticated analysis can be added to the GA3 analysis workflow as in **Figure 6**. For example, instead of measuring the intensity of a complete cilium, differences in intensity within a sub-region of a cilium may be of interest to assess sub-ciliary localization. Differences in intensity within a subregion of a cilium could indicate the protein is accumulating at the tip or the base of the cilium, such as how Gli proteins are enriched at the tip of cilia^48^. In addition, this Ai application can be used to readily identify differences between genotypes or treatment groups. While our lab primarily uses this method to analyze cilia imaged from brain sections or neuronal cultures, it can be applied to images acquired from various cell lines or other tissue types. The flexibility of sample type that this application can be used on makes this method of analysis valuable for many different groups studying primary cilia or any discrete organelle that is being assessed such as mitochondria, nucleus, or ER.

## ACKNOWLEDGMENTS

This work was funded by National Institute of Diabetes and Digestive and Kidney Diseases R01 DK114008 to NFB and the American Heart Association Fellowship Grant #18PRE34020122 to RB. We thank Rich Gruskin General Manager of Nikon Software, Melissa Bentley, Courtney Haycraft and Teresa Mastracci for insightful comments on the manuscript.

