## Supplemental Figures for "Artificial Intelligence Approaches to Assessing Primary Cilia"

**Supplementary Figures:**

**Supplementary Figure 1. Ai training loss graphs**
**A.** and **B.** Graphs showing training loss of segment.ai on neuronal cilia *in vitro* and *in vivo,* respectively.

**Supplementary Figure 2. Ai Assisted Cilia staining intensity measurements of *in vitro* neuronal cilia.**
**A.** and **B.** Representative images of cilia (MCHR1, red) in primary hypothalamic and hippocampal cultures, respectively. A trained Ai in NIS Elements was used to recognize cilia as shown in the binary mask (cyan) and then GA3 was used to measure the intensity of MCHR1 staining in cilia. Distribution of MCHR1 intensity is graphed as percentage of cilia in 1000 A.U. bins for hypothalamic and 2000 A. U. bins for hippocampal cultures. n= 30 cilia in hypothalamic and 106 cilia in hippocampal cultures from 3 animals. Scale bars 10 µm.

**Supplementary Figure 3. General Analysis 3 recipes for cilia analysis**
**A.** Simple General Analysis (GA3) recipe for measurement of cilia length, intensity and Mander's coefficient. **B.** Complex GA3 recipe for measurement of intensity along the length of the cilium using a marker for basal body.

**Supplementary Figure 1**

### A. *In vitro* training

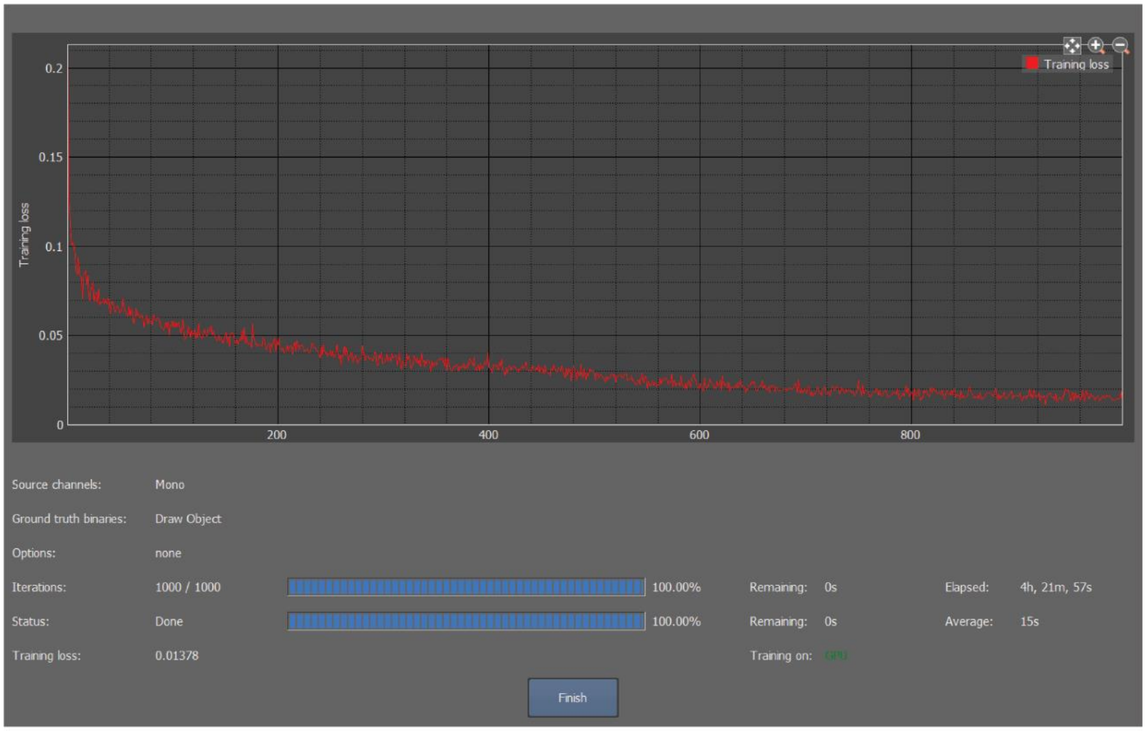


### B. *In vivo* training

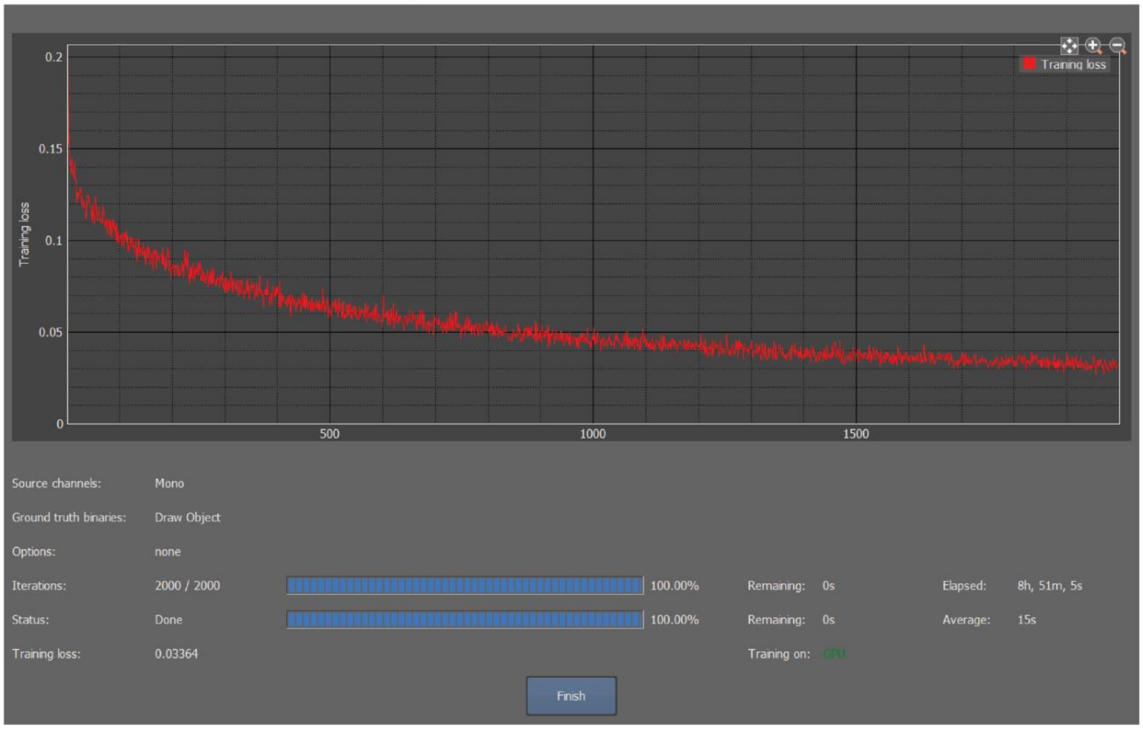

#### Supplementary Figure 2

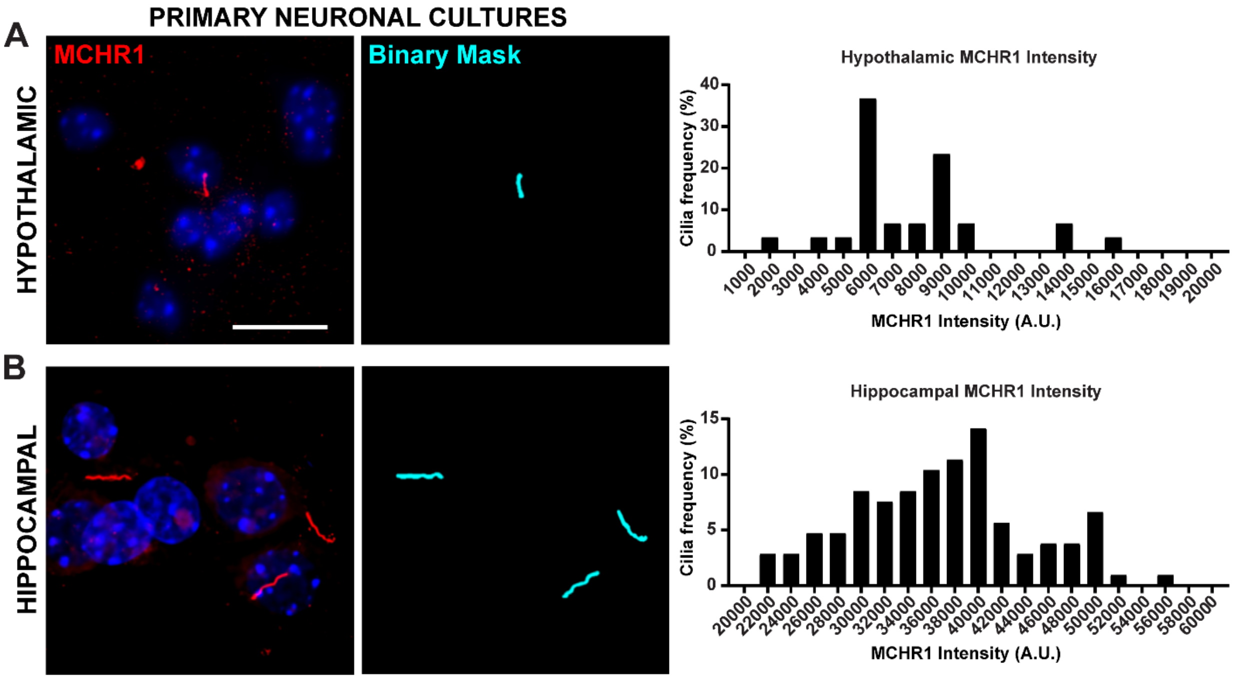

### Supplementary Figure 3

**A** Simple GA3 Recipe (Cilia length, intensity and Colocalization)

**B** Complex GA3 Recipe (Intensity profile along length of cilium)

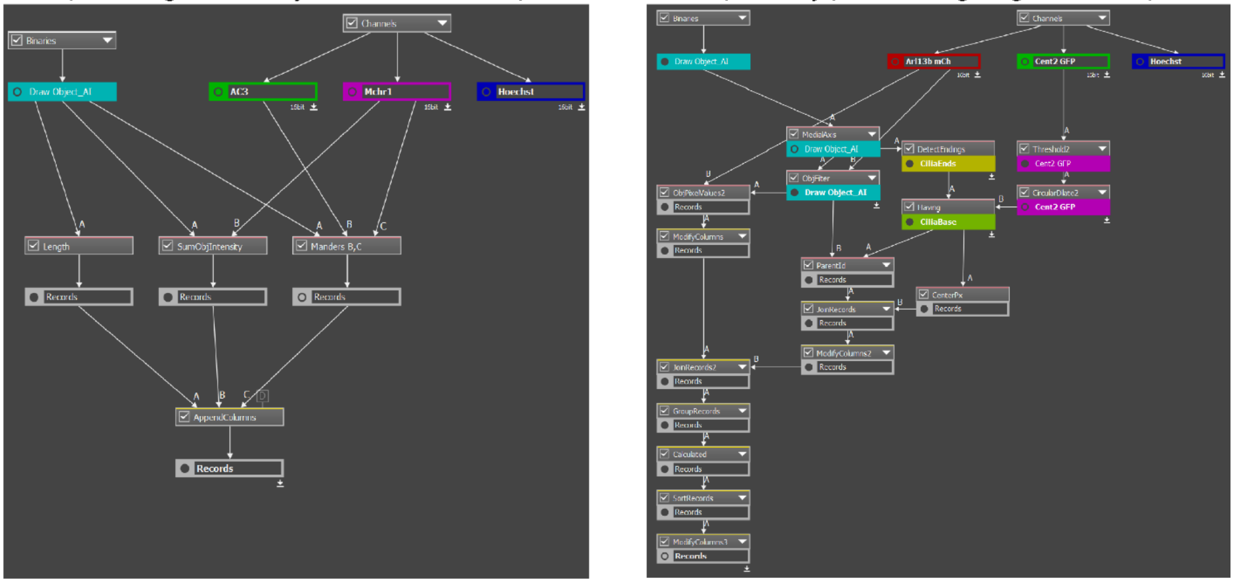
